# Neural rosettes emerge from an interconnected tubular network in human cortical organoids

**DOI:** 10.64898/2026.11.16.767152

**Authors:** Luca Trovò, Gimena Gomez, Ashley Einhorn, Alysson R. Muotri, Lilia M. Iakoucheva

## Abstract

Cortical organoids generate multiple neural rosettes, self-organizing and radially arranged structures that resemble ventricular zones of the developing brain. Here, we show that these rosettes, commonly considered as discrete entities, originate from an interconnected tubular network. We demonstrate that the formation of this continuous structure involves neuroepithelial folding, invagination, and internalization, resembling aspects of neural tube formation. Using fluorescently labeled polyethylene nanoplastics as extracellular tracers, we further show that they accumulate within the developing tubular lumen, consistent with its formation through neuroepithelial folding and internalization. These findings provide new insights into early human brain morphogenesis and suggest the potential utility of cortical organoids for investigating developmental processes relevant to neural tube defects.

## Introduction

Neurodevelopment is a highly regulated process that ensures the proper development of the human brain. The human nervous system arises from the neural tube, a structure formed during early embryonic development. The neural tube itself originates from the neural plate, a specialized region of the ectoderm that undergoes a series of morphological transformations. Through a process known as neurulation, cells of the neural plate proliferate and invaginate, folding into a closed tube that serves as the precursor to the brain and spinal cord (Gilbert 2000; Stiles and Jernigan 2010). Following successive waves of differentiation and structural remodeling, the neural tube gives rise to the major regions of the central nervous system, including the forebrain, midbrain, hindbrain, and spinal cord (Dehay and Huttner 2024; Rakic 2009).

Much of our knowledge of neural tube formation and patterning has historically been derived from animal model studies (Nikolopoulou et al. 2017). However, recent advances in human induced pluripotent stem cell (hiPSC) technologies have enabled the development of brain organoid systems that capture key aspects of human neurodevelopment (Lancaster et al. 2013; Di Lullo and Kriegstein 2017; Rabeling et al. 2024). A defining hallmark of cortical brain organoids is the emergence of neural rosettes (Lancaster et al. 2013; Pasca et al. 2015), radially organized, pseudostratified structures that closely resemble the ventricular zones of the developing human brain (Pasca et al. 2015). These rosettes consist of neuroepithelial progenitor zones, where neural stem cells proliferate, differentiate, and initiate the formation of distinct cortical layers, a process fundamental to mammalian brain development (Chiaradia and Lancaster 2020; Negraes et al. 2021; Trujillo et al. 2019; Fitzgerald et al. 2024).

A recognized limitation of cortical organoids is their tendency to form multiple neural rosettes, which has been thought to reduce their fidelity in modeling human cortical development (Pasca et al. 2015; Chiaradia and Lancaster 2020; Trujillo et al. 2019). To address this limitation, several groups have established protocols for generating single-rosette organoids (Knight et al. 2018; Wang et al. 2022; Kim et al. 2025). These single-rosette-derived organoids are thought to more accurately recapitulate human cortical development by reducing variability associated with multiple rosettes. Yet, despite these advances, the morphogenetic processes that give rise to neural rosettes and their relationship to neural tube formation remain poorly understood.

Despite the widespread use of neural rosettes as morphological hallmarks of cortical organoid development, their three-dimensional developmental origin remains largely unexplored. Typical histological cross-sections and individual optical planes demonstrate that rosettes are spatially separated ventricular-like circular structures, confounding their true architecture. It is not ultimately clear whether they arise independently or emerge through remodeling of a common neuroepithelial architecture. Resolving this relationship requires three-dimensional analysis across developmental stages and visualization of the tubular structures within intact organoids. An independent experimental approach is needed to investigate how extracellular space becomes incorporated into the developing tubular structures.

Here, we show that neural rosettes originate from an interconnected tubular network. The emergence of this continuous structure involves a sequence of neuroepithelial folding, invagination, and internalization that is morphologically reminiscent of neural tube formation, revealing a previously unrecognized parallel between brain organoid morphogenesis and early human brain development. This discovery not only reframes our understanding of organoid architecture but also provides a new model for investigating early neuroepithelial morphogenesis and its potential relevance to neural tube defects.

## Materials and Methods

### iPSCs maintenance

iPSC lines derived from control individuals have been described previously (Fitzgerald et al. 2024; Trujillo et al. 2019; Negraes et al. 2021; Chailangkarn et al. 2016; Adams et al. 2023; Trovo et al. 2026) (**Supplementary Table S1**). iPSC colonies were cultured on Matrigel-coated dishes (Fisher Scientific) with mTeSR Plus medium (STEMCELL Technologies), with medium changes every other day. Colonies were passaged every 5-6 days.

### Generation of cortical organoids

Approximately 1h before initiating organoid differentiation, iPSC colonies were treated with 10 μM ROCK inhibitor (Y-27632; Calbiochem, Sigma-Aldrich). Cells were dissociated using Accutase (Life Technologies) diluted 1:1 in PBS for 10-15 min at 37°C, collected, and centrifuged 200 x g for 3 min. The pellet was resuspended in mTeSR Plus supplemented with 10 μM SB431542 (SB; Stemgent) and 1 μM Dorsomorphin (Dorso; R&D Systems). Approximately 4 × 10^6^ cells were transferred to a single well of a 6-well plate and maintained in suspension under rotation (95 rpm) in the presence of 5 μM ROCK inhibitor for 24 h, allowing the formation of free-floating spheres. Medium was replaced daily.

After 4 days, mTeSR Plus was replaced with Media1 [Neurobasal medium (Life Technologies) supplemented with GlutaMAX, 1% Gem21 NeuroPlex (Gemini Bio-Products), 1% N2 NeuroPlex (Gemini Bio-Products), 1% non-essential amino acids (NEAA; Life Technologies), 1% penicillin/streptomycin (PS; Life Technologies), 10 μM SB, and 1 μM Dorso] for 7 days, with medium changes every other day.

Organoids were then cultured in Media2 [Neurobasal with GlutaMAX, 1% Gem21 NeuroPlex, 1% NEAA, and 1% PS] supplemented with 20 ng/mL FGF2 (Life Technologies) for 7 days, with daily medium changes. This was followed by an additional 7 days in Media 2 supplemented with 20 ng/mL FGF2 and 20 ng/mL EGF (PeproTech), with medium changes every other day.

To promote neuronal maturation, gliogenesis, and network activity, organoids were subsequently transferred to Media3 [Media 2 supplemented with 10 ng/mL BDNF, 10 ng/mL GDNF, 10 ng/mL NT-3 (all from PeproTech), 200 μM L-ascorbic acid, and 1 mM dibutyryl-cAMP (Sigma-Aldrich)]. Beginning at day 28, cortical organoids were maintained in Media 2 for longterm culture, with medium changes every 3-4 days.

### Immunofluorescence

Organoids were harvested at defined developmental time points for immunofluorescence and other analysis. Samples were washed with PBS and fixed in 4% paraformaldehyde (PFA; Thermo Scientific) in PBS at 4°C overnight. Following fixation, organoids were extensively washed in PBS and transferred into a 30% sucrose solution in PBS at 4°C until they sank to the bottom of the tube (typically ≥ 24 h).

For immunostaining, organoids were incubated in blocking solution (PBS supplemented with 2% horse serum and 0.2% Tween-20) for ≥4 h at room temperature, followed by incubation with primary antibodies for 48 h at 37°C on an orbital shaker. After thorough washing, organoids were incubated with secondary antibodies diluted in blocking solution for 24 h. DAPI (1 µg/mL) was added for the last 2 h of incubation to counterstain nuclei.

### Optical clearing and antibodies

For optical clearing following immunostaining and thorough PBS washes, organoids were transferred to RapiClear 1.52 solution (SUNJin Lab). Clearing required 16-24 h for younger organoids and up to 48 h for older samples. Cleared organoids were stored in RapiClear at 4°C until imaging. The following primary antibodies were used: mouse anti-ZO-1 (clone 1/ZO-1; BD Biosciences, Cat# 610967; RRID:AB_398280), rabbit anti-Ki67 (Abcam, Cat# ab15580; RRID:AB_443209) and mouse anti-Nestin (clone 2C1.3A11; Abcam, Cat# ab18102; RRID:AB_444246). The following secondary antibodies were used: goat anti-rabbit IgG Alexa Fluor 488 (Abcam, Cat# ab150077; RRID:AB_2630356) and goat anti-mouse IgG Alexa Fluor 555 (Abcam, Cat# ab150114; RRID:AB_2687594).

### Imaging and 3D reconstruction

Confocal imaging was performed with a Zeiss LSM 880 microscope equipped with FAST Airyscan and 10x objective. For ZO-1 staining, confocal Z-stacks were acquired with a z-step interval of 1.31 µm. Fluorescence filter sets were chosen based on the fluorophores used in each experiment. Images for all conditions were acquired under identical settings, with intensity values monitored during acquisition to avoid signal saturation.

Three-dimensional reconstructions were generated using Imaris software (Bitplane). Image stacks were processed with the automated surface reconstruction Imaris plug-in, applying identical thresholding parameters across all organoids imaged within the same acquisition session.

ZO1 distribution was quantified on 3D confocal stacks of whole cleared organoids. Each organoid was segmented in 3D to obtain its volume and outer surface. For every voxel within the organoid, the distance d to the nearest point of the surface was calculated with a 3D Euclidean distance transform. The maximum distance, *d*_max_, defined the organoid core. To allow comparison between organoids of different sizes, depth was expressed as relative depth (*d/d*_max_), ranging from 0 at the surface to 1 at the core.

Apical ZO1, concentrated at tight junctions and lining the lumens, was separated from diffuse cytoplasmic staining with a 3D white top-hat filter (structuring element of approximately 10 µm). Voxels whose top-hat signal exceeded the organoid median + 6 x median absolute deviation were classified as apical ZO1, isolated objects smaller than 30 µm³ were discarded, and each retained voxel was weighted by its top-hat intensity. For each organoid, the percentage of total apical ZO1 signal in consecutive relative-depth intervals (width 0.1) was calculated, giving a depth profile in which the intervals sum to 100%.

### Lumen morphology classification

Lumen architecture was classified in day-30 organoids that had been cleared, immunostained for ZO1 and imaged as whole-organoid confocal Z-stacks (N = 28 organoids from five iPSC lines). When an image contained more than one organoid, each organoid was segmented and isolated in 3D before analysis. The ZO1-positive apical network of each organoid was then inspected visually in 3D reconstructions and in top and side projections, and each organoid was assigned to one of four categories: “simple lumen”, a single unbranched tubular or ring-like structure; “multiple discrete lumens”, two or more spatially separated luminal structures, none of them branched; “simple branched lumen”, one connected lumen with 1-3 branch points (Y-, T-or E-shaped), or a single tube with separate small rosettes; and “complex branched lumen”, an interconnected network with ≥4 branch points and/or closed loops. Results are reported as the percentage of organoids in each category, for all organoids pooled and for each iPSC line.

### PE nanoplastic characterization and exposure

Fluorescently labeled polyethylene (PE) nanoplastic particles (nominal diameter 50 nm; proprietary fluorescent labeling; LAB621) were used as a marker to trace extracellular material during organoid development. Upon receipt, particle size was confirmed by transmission electron microscopy (TEM; JEOL 1400 plus) following negative staining with uranyl acetate (**Supplementary Figure S1**). Particle size, aggregation state, and stability were further assessed by dynamic light scattering (DLS) using a Zetasizer Nano series instrument (Malvern). PE nanoplastics were suspended at 5 μg/mL in PBS, as a baseline control, and in Media2, to mirror experimental culture conditions. Samples were vortexed and analyzed at room temperature in disposable cuvettes, measuring Z-average hydrodynamic diameter and polydispersity index (PDI) for each condition, followed by zeta potential as an indicator of particle surface charge and colloidal stability. Three technical replicates were processed per sample, and data were analyzed using the manufacturer’s software (Malvern). For experiments, PE nanoplastics were introduced into the culture medium at a final concentration of 5 μg/mL beginning at day 7 of organoid differentiation and maintained through subsequent medium changes until fixation at the indicated time points. This concentration was chosen to approximate the upper range of PE levels reported in human blood (up to ∼4.65 µg/mL;(Leonard et al. 2024)) while providing sufficient fluorescence for imaging.

### Organoid dissociation and caspase-3/7 assay

Apoptosis was assessed on day 30 in organoids continuously exposed to PE (5 µg/mL) or vehicle. As a positive control, unexposed organoids were treated with camptothecin (CPT; Thermo Scientific; 0.01, 0.1 and 1 µM) for 18 h before the assay. For each condition, 10-12 organoids were rinsed twice in PBS with 0.1% BSA and dissociated at 37°C for up to 50 min in HBSS containing Accutase, papain (16.7 U/mL; Worthington) and DNase I (100 U/mL; Worthington), with gentle pipetting every 10 min. Enzymatic activity was quenched with warm Media2, and the resulting cell suspension was filtered through a pre-wet 40 µm strainer and centrifuged at 200 x g for 7 min. Cells were resuspended in cold PBS with 0.1% BSA, counted with a Countess 3 (Thermo Fisher) and plated in triplicate at 3 x 10⁴ cells per well in a 96-well plate (Greiner). An equal volume of Caspase-Glo 3/7 3D Reagent (Promega) was added to each well, followed by shaking at 510 rpm for 30 s and incubated for 30 min. Luminescence was measured using a Spark plate reader (Tecan) as an indicator of caspase-3/7 activity.

## Results

### Neural rosettes form an interconnected tubular architecture in cortical organoids

To investigate the emergence and three-dimensional organization of neural rosettes in cortical organoids, we generated organoids using the differentiation protocol outlined in **Fig. 1A**. Visualization of neural rosettes within organoids presents significant technical challenges. Although rosettes can be detected using conventional optical microscopy, light scattering and the intrinsic thickness of organoids limit resolution (**Fig. 1B**). Immunofluorescence staining of organoid sections followed by imaging provides a more reliable method for visualizing these structures, but this approach restricts analysis to individual slices. However, three-dimensional (3D) reconstruction of intact organoids allows a more comprehensive view of rosette organization. To achieve this, we applied optical clearing (Chung and Deisseroth 2013), which enables whole-organoid clearing and high-resolution imaging without the need for sectioning.

**Figure 1.**
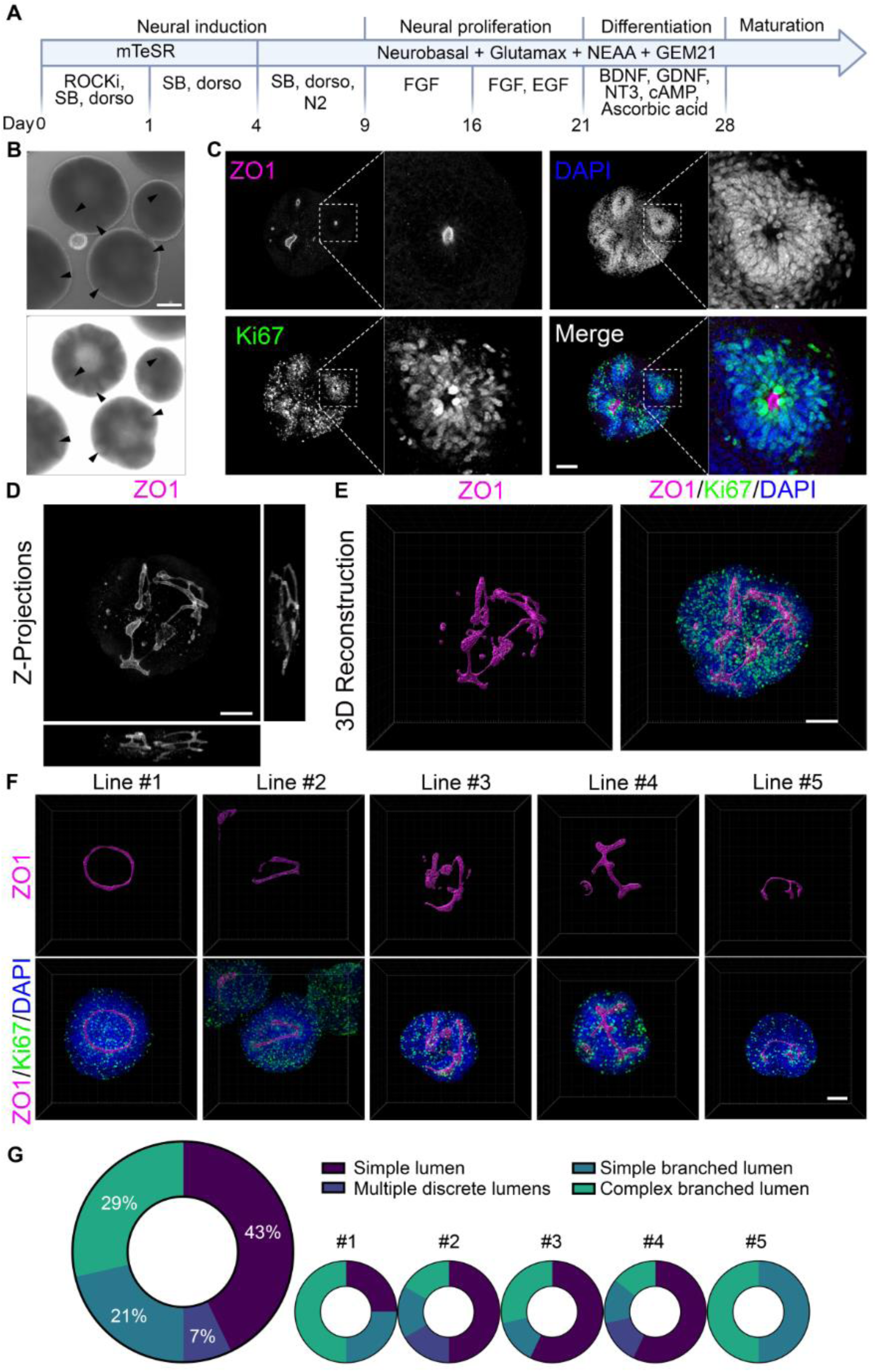
Three-dimensional imaging reveals interconnected tubular architectures in cortical organoids. **(A)** Overview of the cortical organoid differentiation protocol used throughout the study. Human pluripotent stem cells were differentiated into cortical organoids through sequential stages of neural induction, neural proliferation, neuronal differentiation, and maturation. **(B)** Brightfield images of 30-day-old cortical organoids under different illumination settings. *Top*: standard brightfield. *Bottom*: The same organoids imaged with higher illumination, revealing rosettes not visible under standard conditions. Scale bar: 100 μm. **(C)** Confocal images of 30-day-old cleared cortical organoids stained with ZO1 (magenta), Ki67 (green), and DAPI (blue). Scale bar: 100 μm. Insets: Single-plane images showing apparently discrete neural rosettes with proliferative Ki67-positive cells surrounding ZO1-defined centers. **(D)** Z-projection of the ZO1 channel highlighting the continuity of ZO1-positive tubular structures across optical planes. **(E)** 3D reconstruction of a cortical organoid showing tubular organization of interconnected rosettes and the localization of Ki67-positive cells and DAPI-stained nuclei. **(F)** Representative 3D reconstructions of 30-day-old organoids from five iPSC lines. ZO1 staining is shown alone (upper panels) and in combination with Ki67-positive cells and DAPI-stained nuclei (lower panels). Reconstructions illustrate variation in ZO1-positive tubular morphology among the five iPSC lines. Scale bar: 100 µm. **(G)** Distribution of ZO1-positive luminal morphologies in day-30 cortical organoids. Each organoid was assigned to one of four morphological categories based on three-dimensional reconstruction: simple lumen, one unbranched tubular or ring-like structure; multiple discrete lumens, two or more spatially separated luminal structures; simple branched lumen, one connected lumen with 1-3 branch points (Y, T, or E shapes), or a single tube with separate small rosettes; complex branched lumen, a network with ≥4 branch points and/or closed loops. The large donut chart shows the overall distribution, and the smaller charts show the distribution for each iPSC line. N = 28 organoids (Line #1 = 4; Line #2 = 6; Line #3 = 7; Line #4 = 7; Line #5 = 4).

Following immunostaining with DAPI, Ki67 and ZO1, 30-day old organoids were optically cleared (**Fig. 1C**). Here, we define rosettes as radially organized neuroepithelial structures surrounding a central apical lumen, as observed in two-dimensional sections. In single-plane images, rosettes appeared as dense clusters of nuclei (DAPI staining), containing Ki67-positive proliferative cells. ZO1 staining highlighted their apical luminal surface (Elkabetz et al. 2008; Wang et al. 2025), facilitating identification of rosette centers (**Fig. 1C**). Consistent with previous reports (Lancaster et al. 2013; Pasca et al. 2015), these images suggested that rosettes were isolated, radially organized structures. However, Z-projection images of ZO1 staining, which collapse all optical planes into a single view, revealed a strikingly different organization, when seemingly isolated rosettes were interconnected within a continuous tubular network (**Fig. 1D**). We refer to these three-dimensional, ZO1-positive neuroepithelial structures as tubular structures. This interconnected architecture was further validated by 3D reconstruction using IMARIS software, which provided a comprehensive visualization of rosette organization (**Fig. 1E**, **Supplementary Video S1**). To confirm that this tubular architecture was not specific to a single line, we examined multiple organoids across four additional iPSC lines (N=28, range 4-7 organoids per line) for a total of five lines (**Fig. 1F, Supplementary Table S1**). Tubular-like structures were observed in all lines, although their shapes varied (**Fig. 1F**).

To quantify the variability in tubular architecture, we classified the three-dimensional ZO1-positive structures in 28 organoids from five iPSC lines into four morphological categories: “simple lumen”, defined as a single unbranched tubular or ring-like structure; “multiple discrete lumens”, defined as two or more spatially separated luminal structures; “simple branched lumen”, characterized by limited branching; and “complex branched lumen”, characterized by an extensive network with multiple branches and/or closed loops (**Fig. 1G**). Simple lumens were the most frequent morphology (43%), followed by complex branched lumens (29%), simple branched lumens (21%), and multiple discrete lumens (7%). Thus, the vast majority of organoids (93%) exhibited simple or branched tubular architectures. Tubular structures were observed across all five iPSC lines, although the relative frequencies of the four morphological categories varied between lines. These findings demonstrate that simple and branched luminal architectures are common features of cortical organoid development, while also revealing substantial morphological variability among individual organoids.

### Cortical organoids undergo a conserved sequence of neuroepithelial invagination, internalization, and rosette formation

To determine when and how these tubular structures emerge, we imaged organoids at different days of development (**Fig. 2** and **Supplementary Video S2**). We began as early as day 15, when organoids displayed ZO1 localization on the outer surface (**Fig. 2A**). The 3D reconstruction of ZO1 staining revealed that ZO1 covered the entire organoid surface. Additionally, Ki67-positive cells also accumulated on the outer surface, forming what can be described as proliferative neuroepithelium. This structure was apparent even under brightfield microscopy, where organoids around day 15 displayed a darker inner center surrounded by a lighter outer rim-like region (**Fig. 2A**). Immunostaining of the 15-day-old organoids with Nestin, a canonical neuroepithelial marker (Gilyarov 2008), supported the neuroepithelial identity of this layer (**Supplementary Fig. S2A**).

**Figure 2.**
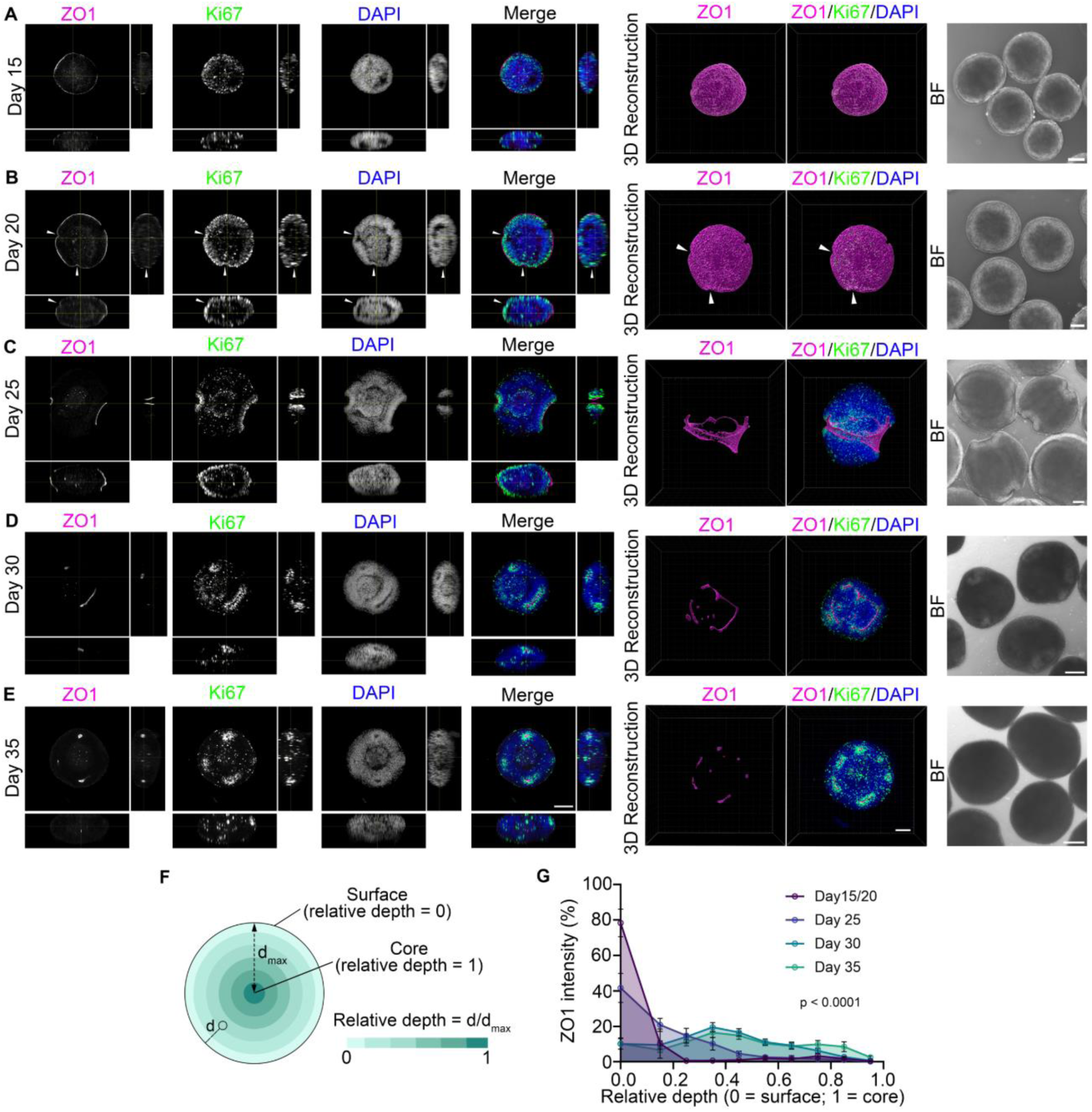
Developmental progression of neuroepithelial folding and tubular internalization in cortical organoids. Orthogonal projection images (left four panels), corresponding 3D reconstructions (next two panels), and brightfield images (right panels) of cortical organoids at different developmental stages (days 15-35). Organoids were fixed, cleared, and stained for ZO1 (magenta), Ki67 (green), and DAPI (blue). Brightfield images show organoids at the corresponding developmental stages. Scale bars: 100 μm (projection images) and 200 μm (reconstruction panels). **(A-B)** At early stages (days 15-20), ZO1 localizes primarily to the outer organoid surface. White arrowheads (**B**) indicate surface indentations consistent with the onset of invaginations. **(C)** By day 25, invaginations expand and wrap around the organoid, producing distinct shapes visible even under brightfield microscopy. **(D)** Around day 30, ZO1 becomes localized internally, forming elongated tubular structures within the organoid. **(E)** By approximately day 35, ZO1-positive structures appear as multiple discrete rosettes, consistent with remodeling of the earlier tubular architecture. **(F)** Schematic of the depth analysis illustrating the calculation of relative depth within an organoid. For each voxel, the distance *d* to the nearest point on the organoid surface was calculated in 3D. The maximum distance, *d*_max_, corresponds to the organoid core. Relative depth was defined as *d*/*d*_max_, ranging from 0 at the surface to 1 at the core. **(G)** Distribution of ZO1 fluorescence as a function of relative depth within the organoid at days 15– 20, 25, 30, and 35. For each depth interval, the graph shows the percentage of the total ZO1 fluorescence signal within that interval. Data are represented as mean ± SEM, with n = 4 (days 15–20), n=12 (day 25), n=28 (day 30), and n=20 (day 35) organoids. Statistical analysis: ZO1 depth distributions were compared across developmental stages using a two-way mixed-effects model (REML; stage x depth bin, organoid as the repeated subject, Geisser-Greenhouse correction), followed by Tukey’s multiple comparisons between stages within each depth bin. The stage x depth interaction was significant (p < 0.0001).

Between days 15 and 20, no major structural changes were apparent, except for the appearance of small indentations on the organoids’ surfaces (**Fig. 2B, white arrows**). By days 20-25, more pronounced morphogenetic changes were observed (**Fig. 2C**). Interestingly, at this stage, organoid shape variations were visible even under the brightfield microscope. Organoids began to lose their perfect circular shape, developing bumps, surface folds and irregularities. ZO1 and Ki67-positive cells became concentrated within specific surface areas, forming small invaginations. These invaginations appeared to wrap around the organoid, creating either a ring-like structure or multiple “arms” that extend around its circumference. The examined organoids displayed invaginations of varying shapes and appearances on their surfaces (**Supplementary Fig. S2B**), suggesting that these morphogenetic events underlie the diversity of tubular-like structures observed at later stages (**Fig. 1F**).

By around day 30, ZO1 staining had shifted from the outer surface to organoid interior, where it was entirely surrounded by proliferative Ki67-positive cells (**Fig. 2D**). At this stage, the tubular structure was localized internally within the organoid, and the invagination pattern likely influenced its shape. As development progressed into day 35, organoids transitioned into a state characterized by discrete, isolated rosettes, consistent with remodeling of the earlier tubular structure (**Fig. 2E**).

It is important to note that due to the inherent variability of organoid maturation (Velasco et al. 2019), not all organoids, even those grown from the same culture, progressed through the same stages of maturation synchronously. At the same time point, some organoids continued to display surface invaginations, while others had already established internalized rosettes. This variability was even more pronounced across different iPSC lines, and thus the developmental days indicated in Figure 2 should be regarded as approximate temporal landmarks rather than absolute benchmarks. Nevertheless, the examined organoids exhibited similar sequence of morphological states, highlighting a conserved developmental trajectory that parallels aspects of neural tube formation.

To quantify these temporal transitions of tubular structure formation, we measured the 3D distribution of ZO1 within whole cleared organoids across developmental stages. For each organoid, the distance of every voxel from the outer surface was calculated in 3D and normalized to the maximum distance (the organoid core), giving a relative depth ranging from 0 (surface) to 1 (core) (**Fig. 2F**). ZO1 was profiled as a function of relative depth (**Fig. 2G**). Between days 15 and 20, ZO1 was almost entirely confined to the organoid surface, with ∼80% of the apical signal located in the outermost depth interval. By day 25, this surface fraction had decreased to ∼40%, and the remaining signal had started to spread into the outer layers of the tissue. By days 30 and 35, only ∼10% of apical ZO1 remained at the surface, while the rest was distributed throughout the organoid interior. The depth distributions differed significantly across developmental stages (stage x depth interaction, p<0.0001). These measurements quantitatively confirm the progressive shift of the apical surface from the outside of the organoid to its interior, and show that the major redistribution of ZO1 toward the organoid interior occurs between days 25 and 30.

### Extracellular PE tracer accumulation supports tubular lumen formation during neuroepithelial morphogenesis

To investigate whether extracellular material becomes incorporated into the developing tubular lumen during neuroepithelial morphogenesis, a process morphologically reminiscent of neural tube formation (Suzuki, Morita, and Ueno 2012; Nikolopoulou et al. 2017), we introduced fluorescently labeled polyethylene (PE) nanoplastic particles (mean diameter ∼50 nm) into the culture medium beginning at day 7, ensuring their presence throughout the period of invagination and tube formation (**Fig. 3A**). Prior to use, PE nanoplastic particles were confirmed to retain their expected size and remain predominantly monodisperse in organoid culture medium, with minimal aggregation (**Supplementary Fig. S1A-B**).

**Figure 3.**
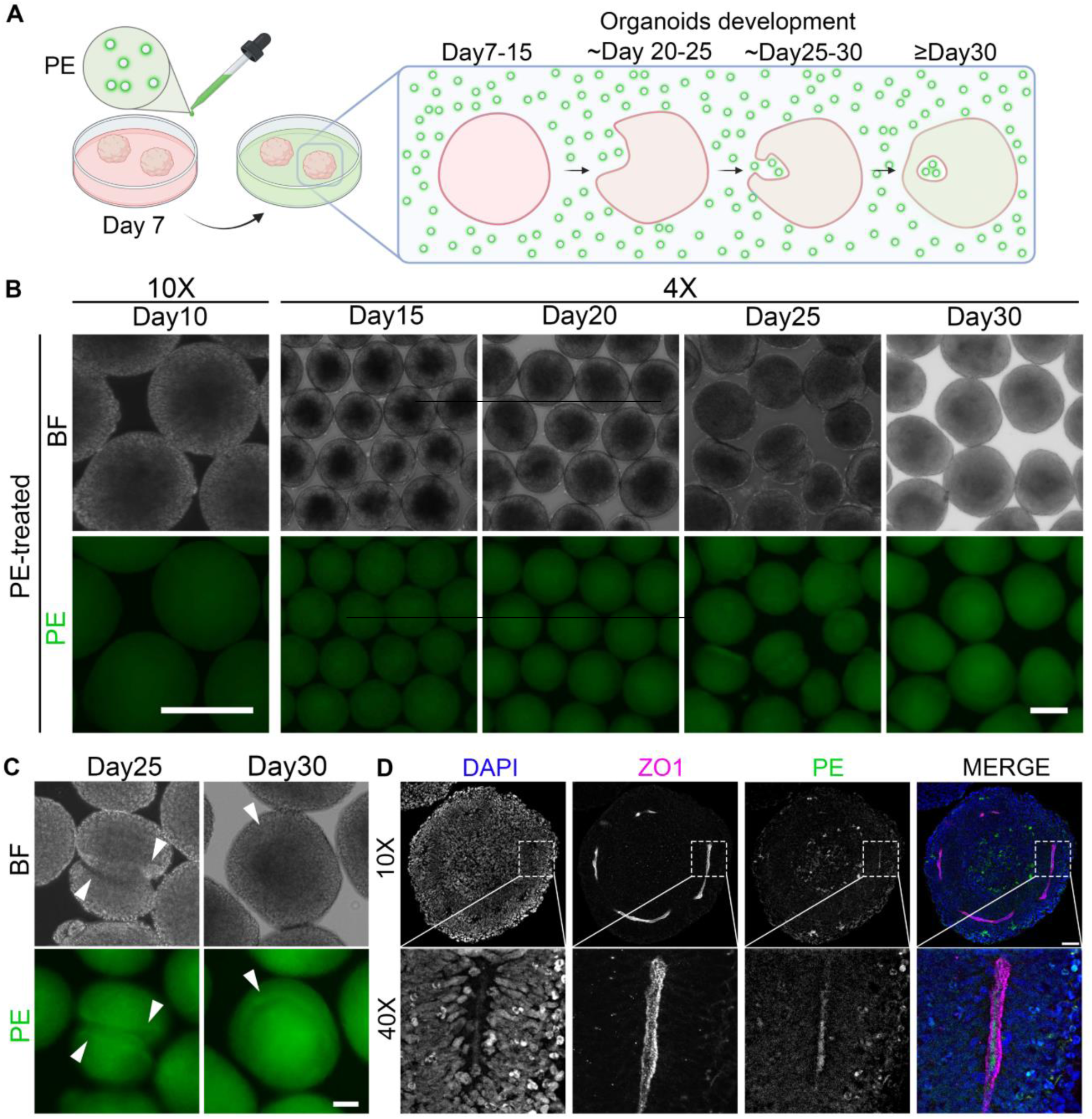
PE nanoplastic particle tracing reveals luminal accumulation during neural tube-like morphogenesis in cortical organoids. **(A)** Schematic representation of the experimental design. Fluorescent PE nanoplastic particles were added to the culture medium beginning at day 7 at each medium change and remained present throughout neural tube-like morphogenesis. As the neuroepithelium invaginates and forms internalized tubular structures, extracellular PE particles become incorporated into the developing lumens. **(B)** Brightfield (upper panels) and PE fluorescence (lower panels) imaging of organoids exposed to PE nanoparticles (5 µg/mL), acquired at five-day intervals from day 10 through day 30. PE nanoparticles were replenished with each medium change. Images were acquired at 10x (day 10) and 4x (days 15–30). Scale bars: 200 µm. **(C)** Brightfield (upper panels) and fluorescence imaging (lower panels) of organoids exposed to 5 µg/ml PE nanoparticles (green) at days 25 and 30. Arrowheads indicate nanoplastic accumulation at sites of invagination (day 25) and within the developing tubular structure (day 30). Scale bar: 100 µm. **(D)** Confocal imaging of a cleared organoid at day 30 stained with DAPI (blue) and ZO1 (magenta), showing PE nanoplastic fluorescence (green) localized within the ZO1-defined lumen. Top row (10x): whole-organoid view; bottom row (40x): magnified inset showing PE signal within the lumen of the tube-like structure. Scale bar: 100 µm.

To determine whether prolonged PE exposure was associated with increased apoptotic activity, we measured caspase-3/7 activity at day 30 following continuous exposure to PE (5 µg/mL). No significant difference was observed between PE-exposed and vehicle-treated organoids, whereas a positive control camptothecin treatment induced a dose-dependent increase in caspase-3/7 activity (**Supplementary Fig. S1C**).

Fluorescence imaging revealed the presence and persistence of PE signal within organoids throughout morphogenesis, with fluorescence detectable from day 10 and persisting through the formation of neural tube-like structures by day 30 (**Fig. 3B**). The fluorescent signal also highlighted the developing tubular architecture, allowing visualization of invaginating structures that were not readily apparent by standard brightfield imaging alone (**Fig. 3C**).

To further investigate spatial relationships between extracellular particles and the developing tubular lumen, organoids were optically cleared and imaged by confocal microscopy at day 30. Fluorescent PE signal was detected within the ZO1-defined lumen, surrounded by the ZO1-positive apical surface. This luminal localization was supported by orthogonal projections and three-dimensional reconstruction (**Fig. 3D**). Quantification of PE fluorescence patterns revealed tubular accumulation in 62-77% of organoids across two iPSC lines (**Supplementary Fig. S3A**). In contrast, PE exposure initiated after rosette formation (day 40) did not result in preferential tubular accumulation (**Supplementary Fig. S3B**). These findings demonstrate that extracellular material present during invagination becomes incorporated into the developing tubular structures, supporting a model of neuroepithelial folding and internalization reminiscent of neural tube formation.

## Discussion

Organoids provide powerful models for studying human neurodevelopment by enabling direct observation of three-dimensional tissue organization and morphogenesis. A defining feature of cortical organoids is the formation of multiple neural rosettes, which are typically viewed as discrete ventricular-like structures and are often considered a source of variability or a limitation of organoid models (Pasca et al. 2015; Chiaradia and Lancaster 2020; Lancaster et al. 2013). Here, we show that neural rosettes do not initially arise as independent structures. Instead, three-dimensional reconstruction across developmental stages reveals that they emerge through remodeling of a continuous neural tube-like architecture that undergoes surface invagination, internalization, and eventual resolution into discrete rosettes.

This developmental trajectory provides new insight into the morphogenetic processes that give rise to neural rosettes. Early-stage organoids form a proliferative neuroepithelial layer with an outward-facing ZO1-positive apical surface. This surface subsequently develops folds and invaginations that extend around the organoid and become internalized, producing a continuous tubular architecture surrounded by proliferating neural progenitors. At later stages, this architecture resolves into the multiple rosettes typically observed in cortical organoids (**Fig. 4**). Thus, structures that appear disconnected in individual histological sections or optical planes can represent different regions of a larger, continuous three-dimensional structure. These findings highlight the importance of examining organoid morphogenesis in three dimensions and across developmental time, rather than inferring developmental relationships from static sections of mature organoids.

**Figure 4.**
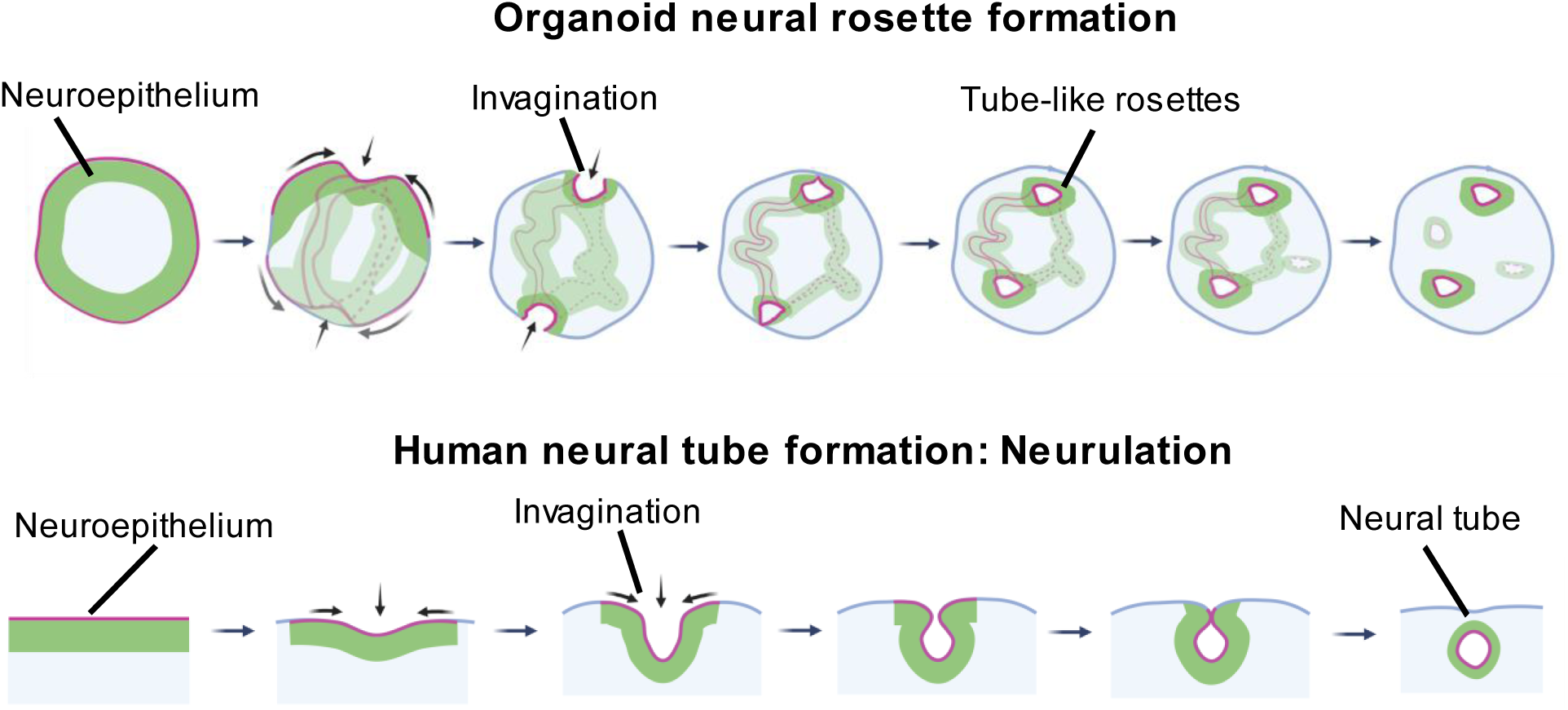
Schematic diagram illustrating morphological parallels between organoid rosette formation and human neurulation. The apical surface is colored in magenta, and the proliferative neuroepithelium in green. Solid colors represent the central plane of the organoid, while lighter shades indicate structures of adjacent planes. Dotted and solid outlines indicate structures in planes below and above the central plane, respectively.

The sequence of morphological transitions observed in cortical organoids parallels key features of primary neurulation, including formation of a polarized neuroepithelium, tissue folding and invagination, internalization of the apical surface, and formation of a lumen-containing tubular structure. The extracellular tracer PE experiment provides additional evidence for the incorporation of extracellular material during this morphogenetic process. PE particles present during the period of invagination became incorporated into the developing tubular structures and were subsequently detected within ZO1-defined lumens **(Fig. 3)**. Although these findings do not establish molecular or mechanistic equivalence between organoid morphogenesis and embryonic neurulation, they demonstrate that cortical organoids undergo a sequence of morphological transformations reminiscent of neural tube formation.

Our findings also change how the presence of multiple rosettes in cortical organoids may be interpreted. Protocols have been developed to generate single-rosette organoids, in part because multiple rosettes are thought to reduce reproducibility and developmental fidelity (Knight et al. 2018; Wang et al. 2022; Kim et al. 2025). Our results suggest that multiple rosettes do not arise independently and that they are not as disorganized as previously thought. To the contrary, they likely emerge through remodeling of a common tubular architecture. Variation in the number, size, and spatial organization of mature rosettes may therefore reflect differences in the geometry of earlier neuroepithelial folding and tubular morphogenesis. Understanding the developmental origins of this variability could provide new criteria for evaluating organoid quality and comparing differentiation protocols.

The transient neural tube-like early stage may also expand the variety of developmental processes that can be investigated using cortical organoids. Most studies use organoids to examine neurogenesis, neuronal differentiation, cortical patterning, and later stages of maturation. Cortical organoids have also been used to investigate the effects of genetic risk variants, including 16p11.2 copy number variants, and environmental exposures, including acetaminophen, on neurodevelopment (Urresti et al. 2021; Trovo et al. 2026). The developmental window identified here provides an experimentally accessible system for investigating earlier morphogenetic events associated with neural tube formation. Organoids could therefore provide a complementary experimental model for studying genetic variants associated with neural tube defects or with neurodevelopmental disorders involving abnormal neuroepithelial morphogenesis (Rabeling et al. 2024). This system may also enable experimental manipulation of signaling pathways, environmental exposures, or candidate preventive interventions during defined stages of neuroepithelial folding and tubular morphogenesis.

The PE nanoplastic particles used in this study primarily served as extracellular tracers to visualize the incorporation of extracellular material into developing tubular structures. However, the ability of nanoscale extracellular material to accumulate within developing lumens raises additional questions about environmental exposures during early neurodevelopment. Nanoplastics have been detected in human placental tissue (Ragusa et al. 2021), and experimental studies suggest that exposure during embryonic development can affect neural tube morphogenesis and neurodevelopment (Huang et al. 2025; Wang et al. 2023; Negraes et al. 2021). Future studies could use the developmental window defined here to investigate how the timing of exposure influences particle accumulation within lumens and whether persistent luminal exposure affects subsequent neural development. These possibilities represent secondary applications of the morphogenetic model established in this study.

Together, our findings reveal a previously unrecognized developmental relationship between the tubular architecture of early cortical organoids and the multiple neural rosettes observed at later stages. Rather than forming as independent structures, neural rosettes emerge through a continuous sequence of neuroepithelial folding, tubular internalization, and structural remodeling. Recognition of this developmental sequence provides a new perspective on rosettes and identifies an experimentally accessible period of neural tube-like morphogenesis. More broadly, our findings illustrate how three-dimensional analysis across developmental time can reveal morphogenetic processes that are obscured by conventional analysis of individual sections or developmental stages.

### Limitations of the study

Although the observed phenomenon was robustly replicated across multiple cell lines in organoids generated using our semi-guided protocol (Fitzgerald et al. 2024), it remains to be determined whether organoids generated using unguided (Lancaster et al. 2013) or fully guided (Pasca et al. 2015) approaches undergo similar developmental transitions. Different differentiation protocols impose distinct signaling environments and tissue geometries that may influence neuroepithelial folding, tubular organization, and rosette formation.

In addition, the developmental sequence described here was reconstructed from fixed organoids collected at successive time points rather than by continuous live imaging of individual organoids. Although the quantitative analyses demonstrate the prevalence of tubular architectures and progressive internalization of the ZO1-positive apical surface, live imaging will be required to directly establish the transitions between surface invagination, tubular internalization, and the subsequent emergence of distinct rosettes. Further studies will also be needed to determine how the timing and geometry of these processes vary among organoids, cell lines, and culture conditions. Finally, because organoids lacking rosettes were excluded during routine quality control, we could not determine the frequency of incomplete or failed tubular internalization.

Finally, our findings demonstrate morphological similarities between the observed organoid process and neural tube formation but do not establish molecular or mechanistic equivalence to embryonic neurulation. ZO1 staining delineates epithelial junctions and luminal boundaries but does not fully characterize apicobasal polarity. Likewise, the localization of extracellular PE particles within developing lumens does not establish the precise mechanism of particle entry or retention. The signaling pathways and cellular mechanisms responsible for neuroepithelial folding and tubular internalization were not investigated here, and perturbation of canonical regulators of neural tube morphogenesis will be required to determine the extent to which the two processes share common mechanisms. Molecular profiling, live imaging, and experimental perturbation of the developmental window identified here will be important next steps toward establishing cortical organoids as a model of early human neural morphogenesis.

## Supplementary Tables, Figures, and Video legends

**Supplementary Table S1.** Donor iPSC lines used for organoid production.

| **Line #** | **Line Name** | **Sex** | **Reference** |
|---|---|---|---|
| 1 | WT83c6 | Male | Trujillo et al., 2021; Fitzgerald et al., 2024 |
| 2 | CVB | Male | Trujillo et al., 2019; Adams et al., 2023 |
| 3 | 121c4 | Female | Negraes et al., 2021 |
| 4 | TD55 | Female | Chailangkarn et al., 2016; Trovo et al., 2026 |
| 5 | TD63 | Female | Chailangkarn et al., 2016; Trovo et al., 2026 |

**Supplementary Table S2.** Source data and statistical analyses underlying quantitative figures.

**Supplementary Figure S1.**
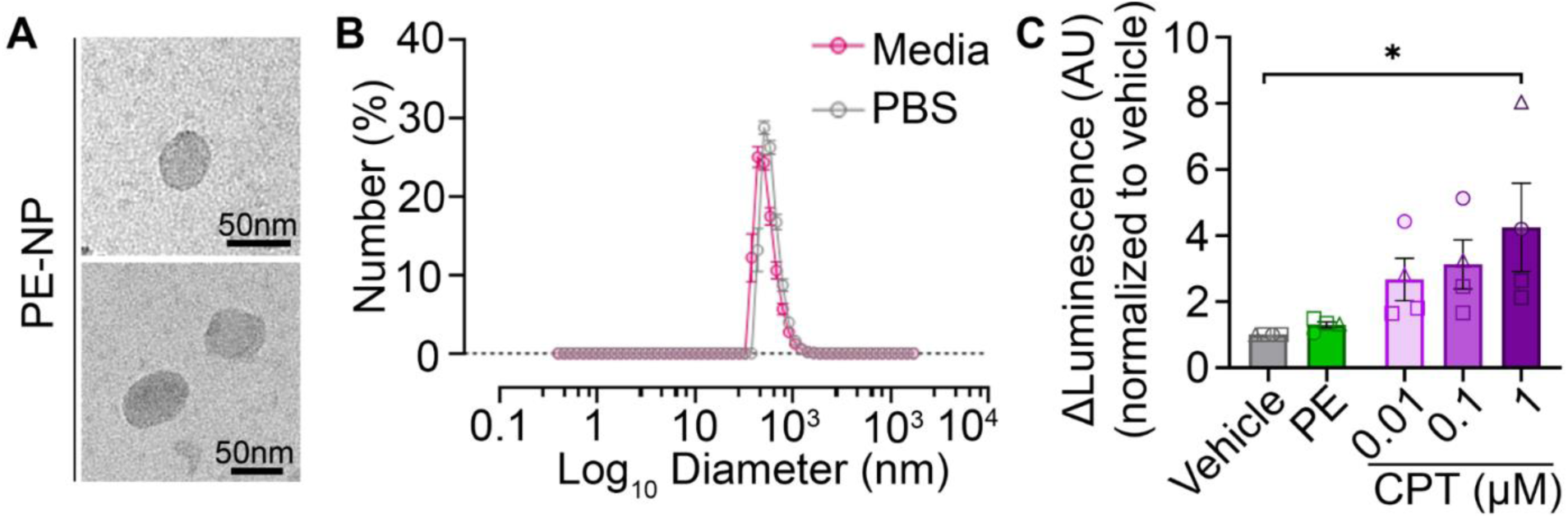
Physicochemical characterization of PE nanoplastic particles and assessment of apoptosis following PE exposure in cortical organoids. **(A)** TEM images of polyethylene (PE) nanoplastic particles (PE-NP), showing particles approximately 50 nm in diameter. Scale bar: 50 nm. **(B)** Number-weighted (Mie-transformed) DLS size distribution of PE nanoplastic particles in organoid culture medium and in PBS, showing a single, overlapping peak in both conditions consistent with a predominantly monodisperse population and minimal aggregation. Data are shown as mean ± SD of three independent experiments, each measured in technical triplicate. **(C)** Quantification of caspase-3/7 activity in day-30 cortical organoids following continuous exposure to PE (5 µg/mL) from day 7, compared with vehicle-treated organoids. No significant difference in caspase-3/7 activity was observed between PE-treated and vehicle-treated organoids. Treatment with camptothecin (CPT) was used as a positive control and resulted in a dose-dependent increase in caspase-3/7 activity. Data are presented as mean ± SEM. Statistical analysis: one-way ANOVA; four iPSC lines across three independent differentiation batches (*p* < 0.05 for the comparison indicated by the bracket).

**Supplementary Figure S2.**
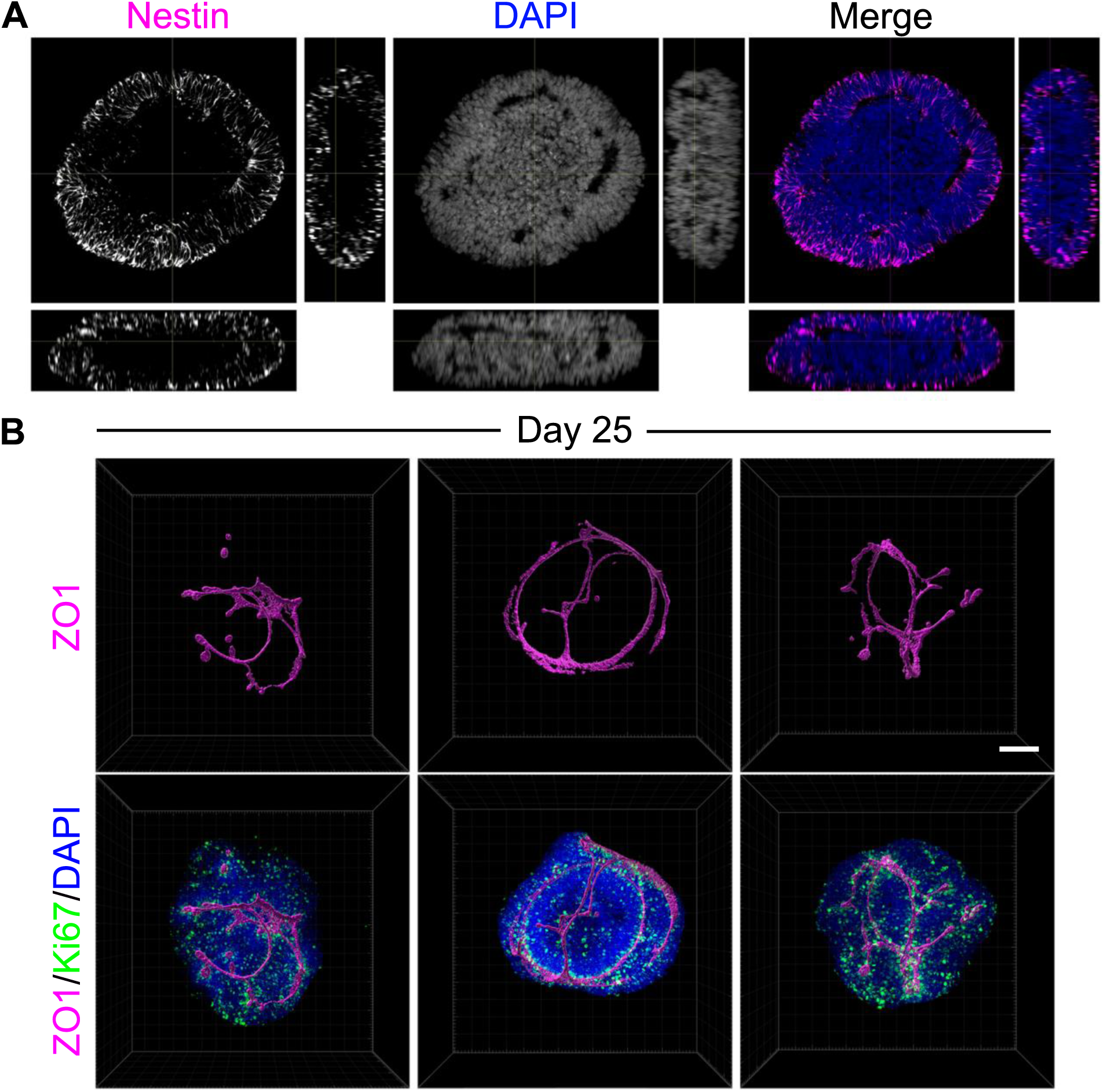
Neuroepithelial development in organoids from day 15 to day 25. **(A)** Orthogonal projections of a 15-day-old organoid immunostained for Nestin (magenta) and counterstained with DAPI (blue), showing a Nestin-positive neuroepithelial layer. **(B)** Representative 3D reconstructions of 25-day-old organoids showing ZO1 staining alone (upper panels) and in combination with Ki67-positive cells and DAPI-stained nuclei (lower panels). Reconstructions highlight variation in the morphology of ZO1-positive epithelial invaginations across organoids during early tubular morphogenesis. Images were obtained from independent differentiation batches distinct from those shown in Fig. 2C, representing a total of three independent batches. Scale bar: 100 µm.

**Supplementary Figure S3.**
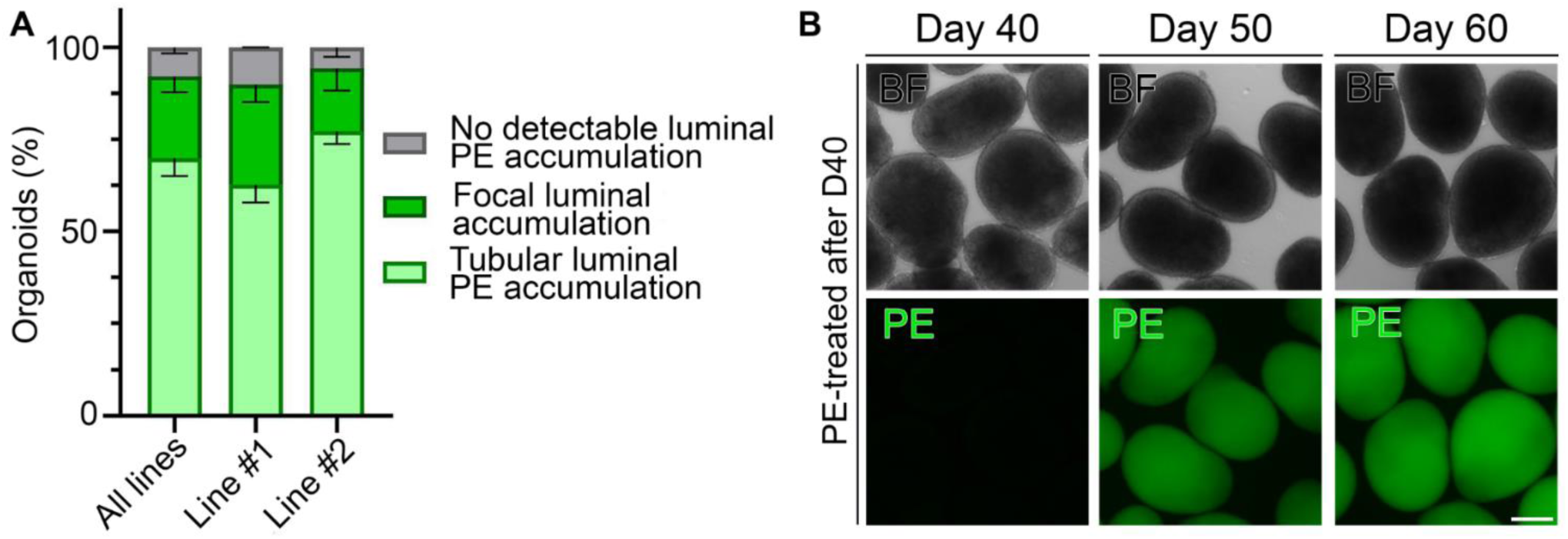
Quantification of PE fluorescence patterns and effects of exposure timing in cortical organoids. **(A)** Quantification of PE fluorescence patterns in cortical organoids exposed to PE beginning at day 7. Percentage of organoids showing tubular PE fluorescence (light green), focal PE fluorescence (dark green), or no detectable preferential PE accumulation (grey), for both lines pooled and for Line #1 and Line #2 separately. Bars show the mean percentage; error bars show SEM. n = 86 (Line #1) and n = 68 (Line #2) organoids from two independent differentiation batches per line. **(B)** Representative brightfield (BF, upper panels) and PE fluorescence (lower panels) images of organoids exposed to PE only after neural rosette formation, with exposure initiated at day 40. Images were acquired at days 40, 50, and 60. No preferential tubular accumulation of PE was observed following late exposure. Scale bar: 100 µm.

**Supplementary Video S1: Neural rosettes form a continuous tubular structure.**

3D reconstruction of a 30-day-old organoid stained with DAPI (blue), ZO1 (magenta), and Ki67 (green), visualized using Imaris software. The video begins with sequential exploration of individual optical planes to illustrate the internal organization of ZO1, followed by a full 3D view of the organoid. The final reconstruction of the ZO1-stained surface highlights an interconnected tubular network, demonstrating that neural rosettes are not isolated entities but part of a continuous tube-like architecture showing neural tube-like morphology.

**Supplementary Video S2: Dynamics of ZO1 distribution during organoid development.**

3D reconstructions of organoids stained with DAPI (blue), ZO1 (magenta), and Ki67 (green) at 15, 20, 25, 30, and 35 days of development. At each stage, the video presents the entire organoid followed by a reconstruction of the ZO1-stained surface. The sequence illustrates the progressive changes in ZO1 localization, initially distributed across the outer surface (days 15 and 20), concentrated within surface invaginations (day 25), reorganized into continuous tubular structures (day 30), and subsequently appearing as discrete rosettes (day 35).

## Acknowledgements

We thank the members of the Iakoucheva Laboratory and Dr. Joseph Gleeson for helpful discussions. This work was supported in part by NIH grants 1R21HD109616 (L.M.I and A.R.M), R01ES033636 (L.M.I and A.R.M), R21MH128827 (L.M.I), and CIRM DISC4-16377 (L.M.I and A.R.M). We also thank the UC San Diego School of Medicine Microscopy Core for access to confocal microscopy facilities and technical support (NINDS grant P30NS047101).

## Author contributions

L.T. and L.M.I. conceived the study, designed the experiments, and wrote the manuscript. L.T. performed the experiments, analyzed the data, and prepared the figures. G.G. contributed to experimental design and manuscript editing. A.E. assisted with organoid culture, imaging, and PE experiments. A.R.M. provided scientific guidance and contributed to manuscript writing. L.M.I. and A.R.M. supervised the project. All authors reviewed and approved the final manuscript.

## Data availability

Full-resolution raw confocal (Airyscan) Z-stacks corresponding to all representative images shown in the figures were deposited to Zenodo as two datasets: Part 1 (DOI: 10.5281/zenodo.23486928) and Part 2 (DOI: 10.5281/zenodo.23510024). Each Z-stack is provided as a multichannel OME-TIFF file with the voxel size embedded in the metadata, and the datasets include a file list linking each image stack to its corresponding figure panel, together with the stains, image dimensions and voxel size. The source data underlying all quantitative graphs, together with the corresponding statistical analyses, are provided in **Supplementary Table S2**. Each worksheet corresponds to an individual figure panel and is labeled accordingly.

